# The haplotype-resolved *Prymnesium parvum* (type B) microalga genome reveals the genetic basis of its fish-killing toxins

**DOI:** 10.1101/2024.03.27.587007

**Authors:** Heiner Kuhl, Jürgen F. H. Strassert, Dora Čertnerová, Elisabeth Varga, Eva Kreuz, Dunja K. Lamatsch, Sven Wuertz, Jan Köhler, Michael T. Monaghan, Matthias Stöck

## Abstract

The catastrophic loss of aquatic life in the Central European Oder River in 2022, caused by a toxic bloom of the haptophyte microalga *Prymnesium parvum* (in a wide sense, s.l.), underscores the need to improve our understanding of the genomic basis of the toxin. Previous morphological, phylogenetic, and genomic studies have revealed cryptic diversity within *P. parvum* s.l. and uncovered three clade-specific (types A, B, C) prymnesin toxins. Here, we used state-of-the-art long-read sequencing and assembled the first haplotype-resolved diploid genome of a *P. parvum* type B, the strain responsible for the Oder disaster. Comparative analyses with type A genomes uncovered a genome-size expansion driven by repetitive elements in type B. We also found conserved chromosomal synteny but divergent evolution in several polyketide synthase (PKS) genes, which are known to underlie toxin production in combination with environmental cues. We identified a specific, approximately 20 kilobase pair comprising deletion in the largest PKS gene of type B that we link to differences of the chemical structure of types A and B prymnesins. Electron-microscopy and flow cytometry confirmed diploidy in the Oder River strain and differences to closely related strains in morphology and ploidy. Our results provide unprecedented resolution of strain diversity in *P. parvum* and a better understanding of the genomic basis of toxin variability in haptophytes. The reference-quality genome will help to understand changes to microbial diversity in the face of increasing environmental pressures, and provides a basis for strain-level monitoring of invasive *Prymnesium* in the future.

## Introduction

In the summer of 2022, an anthropogenic environmental disaster struck the Central European Oder River, resulting in a significant loss of aquatic life due to the proliferation of a strain of the toxin-producing microalga, *Prymnesium parvum* s.l. (‘*sensu lato*’ - in a wide sense) (Haptophyta, Prymnesiophyceae, Prymnesiales, Prymnesiaceae).^1^ The toxins released by this brackish-water mixotroph, which measures only 5–10 micrometers and carries two flagella for active movement and a specialized organelle (haptonema) for attaching to prey, caused a catastrophe, leading to the death of a thousand metric tons of fish, mussels, and snails along the entire Oder River in Poland and Germany.^1–3^ The invasive *Prymnesium* as the cause of massive fish kills has been identified since the mid 20^th^ century,^4,5^ but only in the last three decades has much of its cryptic diversity been recognized. Electron-microscopy revealed variation in the organic scales among strains or between stages of what may be a haplo-diplontic life cycle.^6–8^ Phylogenetic analyses revealed two (ITS1)^7^ or three (ITS1 or ITS1+2)^1,9–11^ different clades (types A, B, C)^10^ shown by evolutionarily relatively conserved DNA markers. Later on, toxicological analyses revealed clade-specific allelopatic toxins (prymnesins), corresponding to these three clades.^10^ These compounds consist of a ladder-frame polyether backbone, whose length defines the type, as well as various pentose and/or hexose units.^10,12^ More recently, flow-cytometry, transcriptomics,^13,14^ and whole genome sequencing^15,16^ indicate that *P. parvum* s.l. is a complex of at least 40 genetically distinct strains (i.e., forms within types A, B, C) that differ in genome size and/or ploidy and produce type-specific prymnesins and even strain-specific mixtures of differently glycosylated/halogenated prymnesin variants.^15,16^ The polyketide-like chemical nature of these prymnesins suggests that polyketide synthases (PKSs) contribute to their biosynthesis,^14,17^ while glycosyltransferases are essential for polysaccharide modifications.

Thus, to uncover the hidden diversity of *P. parvum* s.l., unambiguous identification of these harmful bloom-forming algae requires genomic data.^16^ This will also facilitate the identification of the prymnesin types, which is of particular interest as not only its structure but also its toxicity is type-dependent.^18^ To understand how the genes of the major protein families enable strain-specific toxin production and evolution, comparative genomics involving additional chromosome-scale high-quality reference genomes of *P. parvum* are essential. This will contribute to tackling the future impact of this globally relevant threat, which is of increasing relevance due to freshwater salinization and climate change.^1^

Here, we present the haplotype-resolved diploid genome of *P. parvum* type B (ODER1), the strain that caused the Oder disaster. Compared to a diploid type A genome, we show enormous, though evenly-distributed size expansion of all ODER1 strain chromosomes. We also find ploidy shifts in the most closely-related B-type genome from Denmark that turns out to be triploid. Analysis of PKS genes shows evolutionary changes between type A and B, and — in case of glycosyltransferases — even between closely related B-types that may explain different toxin structures and potential ecologically relevant toxicity.

## Material and Methods

### Sampling and cultivation of Prymnesium parvum ODER1

A water sample from the Oder River was taken in Kostrzyn (river-km 617) on the 19^th^ of August 2022. The sample was then filtered (5 μm) and the retained cells transferred to water originating from Lake Müggelsee in Berlin that was sterile-filtered, autoclaved, and supplemented with NaCl to reach a concentration of 0.2%. To avoid mixing of different substrains that may have existed and to minimize contamination, we started from single *P. parvum* cells that were isolated using a micromanipulator (MMO-202ND; Narishige) equipped with a microinjector (CellTramm Oil; Eppendorf) that held a capillary (Figure 1A). This cell-line was named ‘ODER1’ (used hereafter). Isolated cells were propagated in f/2 Medium (Culture Collection of Algae, University of Göttingen, Germany; https://sagdb.uni-goettingen.de/culture_media/) at a salinity of 0.5%, 20 °C, and a 12 h:12 h light-dark cycle (80 μmol photons m^−2^ s^−1^). To quickly obtain high cell densities needed for RNA sequencing (see below), some cells were separated and propagated in 1/2 SWES Brackish Water (Culture Collection of Algae, University of Göttingen, Germany) at a salinity of 3%.

**Figure 1.**
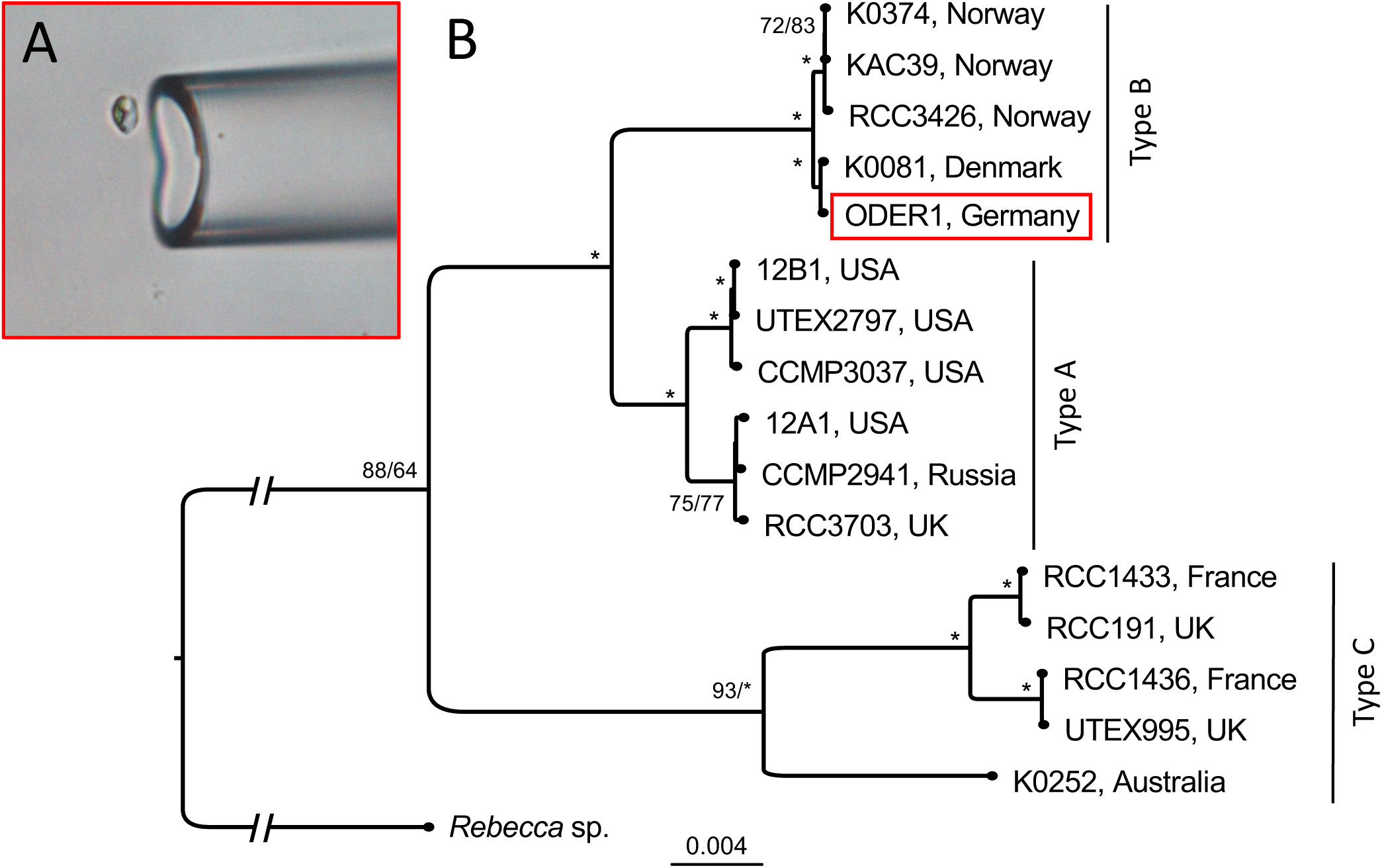
Strain isolation and phylogenetic relationship of *Prymnesium parvum* ODER1. (A) Photomicrograph showing the isolation of a living cell of the ODER1 strain with a micromanipulator; from a single cell, all material used for whole genome sequencing was raised. (B) Maximum likelihood phylogeny of 16 *P. parvum* s.l. strains based on chloroplast genomes, rooted with the haptophyte *Rebecca* sp. Node labels indicate UFBS/SH-aLRT support values. Values >95% are indicated by asterisks. Reference-quality genomes exist only for the type B clade that we newly report here (red box) and for the type A clade (12B1, UTEX2797, CCMP3037).^15,16^ The closest relative of the diploid *P. parvum* ODER1 strain, K-0081, has a triploid genome according to our SNP allele frequency analysis and flow cytometry data (Figure 4).

### High Molecular Weight (HMW) DNA extraction and sequencing

Once *P. parvum* ODER1 cell concentrations reached approximately 1 Mio. cells mL^-1^, ca. 400 mL of culture were isolated following the nuclei isolation protocol published at www.protocols.io by Auber and Wisecaver.^19^ High molecular weight (HMW) DNA was extracted from isolated nuclei using the Nanobind plant nuclei kit (Pacific Biosciences) according to the manufacturer’s protocol. DNA quantification was conducted using UV-spectroscopy (Nanodrop) and Fluorometry (Qbit). To check DNA fragment size quality, a sequencing run was performed on a MinIon sequencer (Oxford Nanopore Technologies (ONT, Oxford, UK) using 1 µg HMW DNA (sheared five times by a G23 needle), the LSK-110 library preparation kit, and a R9.4.1 flow cell. ONT sequencing reads, basecalled by Guppy 6 and the dna_r9.4.1_450bps_plant_sup.cfg model, passed N50 read-length >15 kbp. The final sequencing of the HMW DNA was done on a Pacbio Revio sequencer at Novogene (UK) using circular consensus read mode (CCS/HIFI).

### Prymnesium parvum phylogeny based on chloroplast DNA

ONT MinIon data obtained from test sequencing *P. parvum* ODER1 DNA extractions was assembled using WTDBG2^20^ and polished using the ONT-long-reads by FLYE.^21^ This yielded a single contig chloroplast genome. Similarly, an outgroup chloroplast genome was produced from a sample of *Rebecca* sp. (strain ID 3408, CCAC, University Duisburg-Essen), a haptophyte from the Pavlovaceae family. Short read genome assemblies of other *P. parvum* genomes from Wisecaver et al.^16^ were downloaded from the corresponding figshare repository and aligned to the ODER1 chloroplast genome by Minimap2.^22^ Alignments were converted to maf format (paftools.js view -f maf) and screened for ortholog matches using last-split.^23^ Pair-wise maf files were combined into a multiple alignment using MULTIZ.^24^ The multiple alignment maf file of 17 taxa was converted to aligned fasta format, nucleotide residues with fewer than 15 aligned taxa were removed. A maximum likelihood phylogenetic tree was calculated with IQ-TREE 2.^25^ The best-fit model (K3Pu+F+I+G4) was chosen using the Bayesian inference criterion (BIC) in IQ-TREE 2. Ultra-fast-bootstrap (UFBS)^26^ and Shimodaira–Hasegawa-like approximate likelihood ratio test (SH-aLRT)^27^ methods were applied to calculate branch support values.

### Hi-C library construction and sequencing

The Arima High Coverage HiC Kit (Arima Genomics, Carlsbad, CA, USA) was used to construct a Hi-C sequencing library. We used about 7.5 × 10^7^ *P. parvum* cells and followed the manufacturer’s protocols for nucleated blood. The sequencing library was constructed using the ARIMA protocols for the Accel NGS 2S Plus DNA Library Kit (Swift Biosciences). This library was amplified by nine cycles of PCR. As a quality check, the library was sequenced on our in-house MinIon device as described above. The production-scale sequencing of the Hi-C library was performed on a NextSeq2000 sequencer (Illumina) using 150 bp paired-end read mode at the Berlin Center for Genomics in Biodiversity Research.

### Diploid genome assembly

The longest 1.5 million CCS/HIFI reads (N50 read length 19,300 bp; 29.4 Gbp in total) were used for *de novo* assembly with hifiasm^28^ (0.19.6-r595). Hi-C Illumina data were included to support improved haplotype phasing (options: --h1 --h2). ONT reads were included to allow for gap closure (option: --ul). The Hi-C Illumina reads were independently mapped to the resulting hifiasm haplotype 1 or haplotype 2 contigs using chromap,^29^ and these were scaffolded to chromosome-scale by YaHS.^30^ The Hi-C scaffolds were manually curated using juicebox,^31^ and bacterial contamination could be removed as these contigs had clearly reduced Hi-C signals. A few gaps in the assemblies could be closed as neighboring contigs had long overlaps. The two haploid assemblies were aligned to each other with Minimap2 and stringent mapping parameters “-x asm5” for genomes with divergence of less than 5%,^22,32^ results were plotted in a dotplot fashion using minidot.^33^ The haploid assemblies were also compared to the *P. parvum* 12B1 type A reference genome^16^ (Minimap2, minidot but using less stringent alignment parameters: -x map-ont). A genome browser for the ODER1 assembly has been established at http://genomes.igb-berlin.de:8081/ (will be made publicly available upon acceptance of the manuscript).

### RNA extraction, sequencing, and transcriptome assembly

Cultured *P. parvum* strains UTEX2797 and RCC1436 were obtained from the University of Copenhagen and raised at the growth conditions described above for ODER1 (3% salinity). After attaining cell densities of about 1 million cells/mL, 130 mL of each culture (ODER1, UTEX2797, RCC1436) were filtered using GF/F glass fiber filters (pore size 0.7 µm), and total RNA was extracted using the Transcriptome RNeasy PowerWater Kit (Qiagen, Germany) following the manufacturer’s instructions. Libraries were prepared with the TruSeq stranded mRNA library protocol (poly A selection). Transcriptomes were then sequenced (PE 150 bp) at Macrogen Europe on the Illumina NovaSeq platform. Reads were trimmed with Trimmomatic v. 0.39^34^ (using the options LEADING: 3, TRAILING: 3, SLIDINGWINDOW: 4:15, and MINLEN: 36) assembled with Trinity v. 2.10.0 and proteins were predicted with TransDecoder v. 5.5.0^35^ using default settings, respectively.

### Repeat analysis and comparison

*De novo* repeat analysis was performed on *P. parvum* ODER1 as well as on *P. parvum* 12B1 by RepeatModeler/RepeatMasker.^36^ Repeat annotations of different repeat classes were summarized by the script “buildSummary.pl” to allow for comparison between the different genomes.

### Annotation

A set of transdecoder proteins from our *P. parvum* types A, B, and C transcriptome assemblies (UTEX2797, ODER1, RCC1436, respectively) and Wisecaver et al.^16^ (strain 12B1 annotated proteins) was compiled, and splice-aligned with the genome assembly using miniprot^37^ with gtf output. Transcript sequences were splice-aligned with the genome assembly using Minimap2 (-x splice), supported by a splice junction file, generated from the prior protein alignments. Minimap2 sam output was converted to gtf format. The strand of the mRNA alignments in the gtf file was corrected using the information in the sam “ts:” fields, if necessary. The resulting gtf files of genomic exon coordinates from protein and transcript alignments were combined using Stringtie^38^ and TACO,^39^ and the genomic coordinates of CDS exons were calculated by transdecoder.^35^ All resulting gene-models were functionally annotated by EggNog,^40^ best protein matches (LAST aligner)^41^ and BUSCO.^42^ A single best gene model was chosen from a cluster of gene models according to scoring of its functional annotation or its CDS length (if no functional annotation assigned).

Some genes coding for polyketide synthases (PKSs) were difficult to annotate, because they are extremely large, are only weakly expressed and no reference proteins were available. We found that these genes could be reasonably well annotated by Genscan^43^ *ab initio* gene prediction, while other gene prediction tools like Augustus failed. Thus, we performed Genscan prediction on several *P. parvum* genomes to be able to perform a PKS gene family analysis.

### Estimation of DNA content by flow cytometry and ploidy level assignment using genomics

*Prymnesium parvum* strain K-0081 was obtained from the Norwegian Culture Collection of Algae (NORCCA, Oslo) and propagated in f/2 medium at a salinity of 1%. The DNA content of *P*. *parvum* ODER1 and its closest relative *P*. *parvum* K-0081 was estimated using propidium iodide flow cytometry (PI FCM). To do so, 1 mL of well-grown culture was centrifuged (5 min, 2040 *g*; Eppendorf) and the superfluous medium was removed by pipetting. The cell pellet was flash-frozen in liquid nitrogen, causing a rupture of cells and release of the nuclei. Next, 350 μL of ice-cold nuclei isolation buffer LB01 (15 mM Tris, 2 mM Na_2_EDTA, 0.5 mM spermine tetrahydrochloride, 80 mM KCl, 20 mM NaCl, 0.1% (v/v) Triton X-100; pH = 8.0)^44^ was added. The resulting suspension was thoroughly shaken and kept on ice. Nuclei of two flowering plant species, selected to closely match the DNA content of the investigated sample without overlapping, *Solanum pseudocapsicum* (2C = 2.59 pg)^45^ and *Carex acutiformis* (2C = 0.82 pg)^46^ were used as internal standards for strain K-0081 and ODER1, respectively. To release the nuclei of the standard, a ca. 20 mg piece of fresh leaf tissue was chopped with a razor blade in a plastic Petri dish with 250 μL of ice-cold LB01 buffer. Both suspensions (algal and standard nuclei) were mixed thoroughly and filtered through a 42 μm nylon mesh into a 3.5 mL cuvette fitting the flow cytometer. Following 20 min incubation at room temperature, a staining solution consisting of 550 μL of LB01 lysis buffer, 50 μg mL^−1^ propidium iodide, 50 μg mL^−1^ RNase IIA and 2 μL mL^−1^ β- mercaptoethanol was added. After 5 min incubation at room temperature, the relative fluorescence of at least 10,000 particles was recorded using a CytoFLEX S cytometer (Beckman Coulter, Indianapolis, IN, USA), equipped with a yellow-green laser (561 nm, 30 mW). Histograms were analyzed using CytExpert 2.4.0.28 software (Beckman Coulter). The DNA content was calculated as sample G_1_ peak mean fluorescence/standard G_1_ peak mean fluorescence × standard 2C DNA content.^47^ To minimize random instrumental shift, both strains were analyzed at least three times on separate days and the measurements averaged. To corroborate DNA content differences, simultaneous analysis of strains ODER1 and K-0081 was performed.

We analyzed the allele frequency distribution of variants in K-0081 and ODER1 genomes. The HIFI data of *P. parvum* ODER1 were mapped to the ODER1 haplotype 2 assembly using Minimap2 (-a -x map-hifi), Illumina data for *P. parvum* K-0081 was mapped using parameters for short reads (-a -x sr). Samtools served to create sorted bam files of the data. CLAIR3^48^ was applied to call variants on both datasets. From the resulting vcf files, we calculated the percentage of alternate allelic reads for each variant with total read coverage >=20 and plotted this as a frequency distribution.

### Transmission electron microscopy

Whole-mount preparations of *P. parvum* cells (ODER1 and K-0081) were used for transmission electron microscopy (TEM). The cells were fixed for 1 min in 2% OsO_4_ vapor, rinsed in distilled water, and subsequently stained with 2% aqueous uranyl acetate (1–3 min). The cells were then rinsed one more time and air-dried prior to examination using a Philips CM 120 BioTwin electron microscope. Micrographs were edited with Photopea (www.photopea.com) to mask tiny holes in the formvar film coating the TEM grids.

### qPCR-assay to quantify Prymnesium parvum in water samples

Samples were taken monthly from March 2023 until February 2024 and fixed with Lugol solution. As a gold standard, samples from the Oder catastrophe (n = 80) in summer 2022 were counted microscopically (10x 100) to calculate the correlation factor between ITS1 copy number and *Prymnesium* cell counts. Each 10 mL sample was centrifuged at 4600 *g* [4 °C, 1 h 45 min] and resuspended in 360 µL of buffer ATL (Qiagen). DNA was extracted in duplicate with the QIAamp DNA Mini Kit (Qiagen) followed by cleanup with the OneStep PCR Inhibitor Removal Kit (Zymo Research). qPCR quantification was carried out in duplicate for each replicate with primers PrymF2239 (5’-CACATCCGATCGTGTCTGC-3’) and PrymR2384 (5’-GCACAACGACTTGGTAGG-3’) at 96 °C (3 min), followed by 40 cycles of denaturation at 96 °C (30 s), annealing at 67 °C (30 s), extension at 72 °C (30 sec) [25 µL: 2 U Platinum™ Taq DNA-polymerase, 4 µL extracted DNA, 1.5 mM MgCl_2_, 9.6 pg µL^-1^ BSA, 400 nM each primer, 0.2 mM dNTPs, 1x SYBR Green I] and calculated from a standard dilution series of a PCR product (quantified with QuantiFluor^®^ dsDNA System, Promega with a DeNovix DS-11 spectrometer, Biozym). Cell equivalents were calculated from the ITS1 copy number (R^2^ = 0.87).

## Results

### Chloroplast phylogenetic tree assigns Prymnesium parvum ODER1 to the type B clade

Using our long-read test data (ONT MinIon), we assembled a complete chloroplast genome of *P. parvum* strain ODER1. Screening short-read assemblies^16^ of other *P. parvum* s.l. strains for chloroplast sequences enabled us to calculate a well supported phylogenetic tree of type A, B, C strains (Figure 1B). *Prymnesium parvum* ODER1 is most closely related to a type B strain isolated in 1985 from brackish water in northwestern Denmark (K-0081) and forms a well supported clade with other type B strains from Norway (RCC3426, KAC-39, and K-0374), which suggests that their similarity may be explained by geographic proximity. This corroborates prior results from ITS sequencing of ODER1.^1^

### Prymnesium parvum ODER1 diploid genome assembly and comparison to type A assemblies

Using state-of-the-art HIFI sequencing and Hi-C data (Figure 2), we obtained the first diploid (2n) genome assembly of a *P. parvum* strain, making it the first reference quality assembly of a type B strain. Compared to publicly available type A reference assemblies,^15,16^ statistics and BUSCO scores (Tables 1 and 2) are consistently improved in our assembly. The most striking difference is the genome size. The sizes of our two haploid *P. parvum* ODER1 assemblies (236/237 Mbp) exceeded those of A-type strains 12B1 (94 Mbp) and CCMP3037 (107 Mbp) by a factor of more than two. We did not compare our ODER1 assembly to type A UTEX2797, because two publications present disagreeing assembly results for this strain,^15,16^ which may be due to high heterozygosity, hybridization or issues of cultivation.

**Figure 2.**
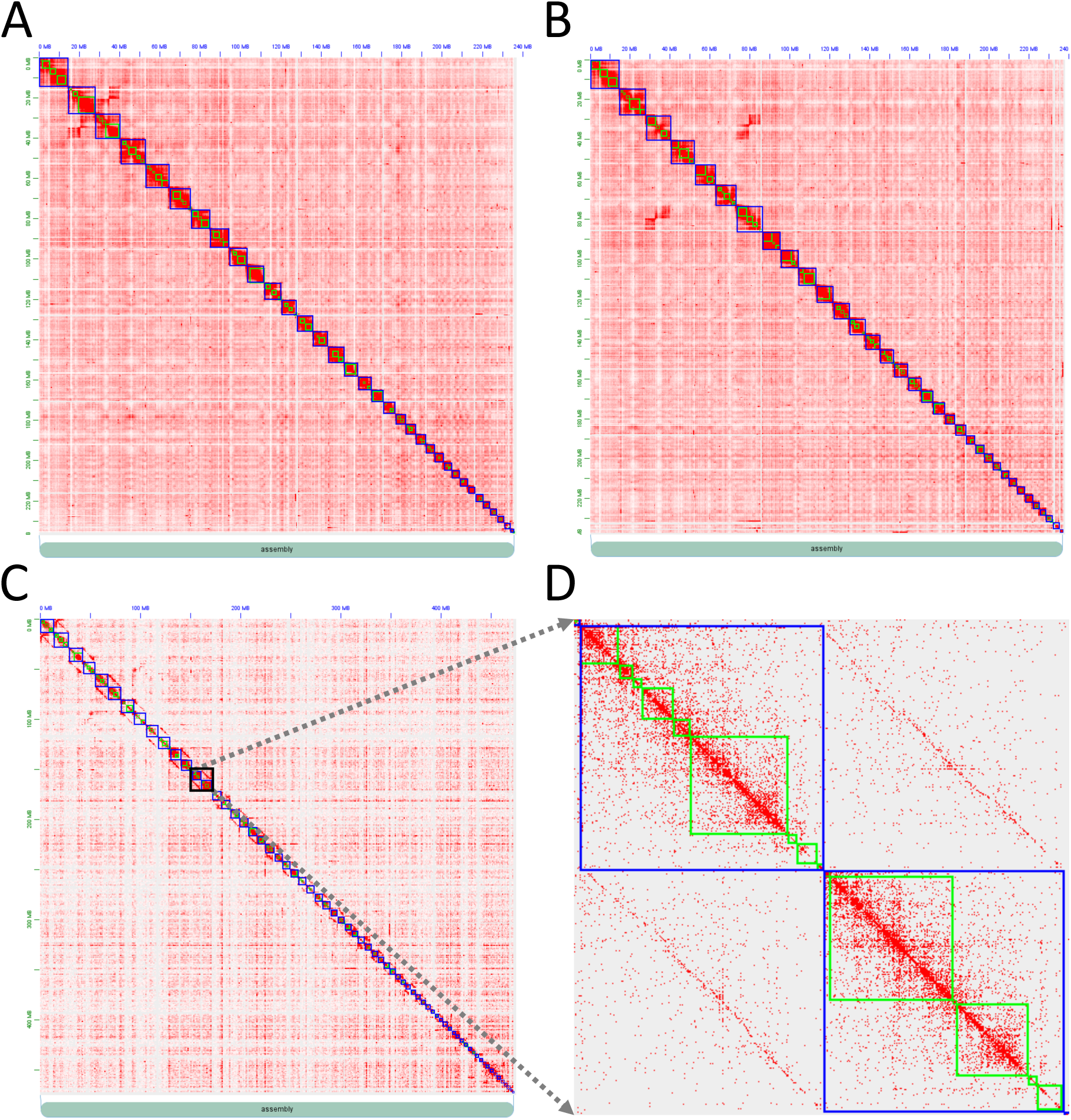
Hi-C interaction matrices of the two *Prymnesium parvum* ODER1 haplotypes. (A) Hi-C map of haplotype 1 (1n = 34) of *P. parvum* ODER1. (B) Hi-C map of haplotype 2 (1n = 34) of *P. parvum* ODER1. (C) Combining both haplotypes (2n = 68) in a single HiC-map shows (D) typical patterns of a diploid assembly (slight signal between haplotypes, while stronger signal within haplotypes).

**Table 1.** Diploid long-read genome assembly results of *Prymnesium parvum* ODER1. The diploid genome could be resolved into two chromosome-level haploid genome assemblies of highly similar quality (hap1, hap2) using HIFI, ONT, and Hi-C sequencing data. Comparison with *P. parvum* type A genomes shows highly increased genome size and repeat content in type B *P. parvum* ODER1, despite diploidy of both strains.

|  | *P. parvum* ODER1<br>(this study) |  | *P. parvum* 12B1<br>(Wisecaver et al.)<sup>16</sup> | *P. parvum* CCMP3037<br>(Jian et al.)<sup>15</sup> |
| --- | --- | --- | --- | --- |
| Type | B | B | A | A |
| Haplotype | hap1 | hap2 | collapsed | collapsed |
| Total length [bp] | 236,176,151 | 237,246,831 | 93,538,114 | 107,321,770 |
| Chromosomal scaffolds | 34 | 34 | 34 | 34 |
| Contig number | 314 | 288 | 225 | 362 |
| Scaffold n50 [bp] | 8,642,580 | 8,552,854 | 3,203,049 | 3,786,890 |
| Contig n50 [bp] | 1,791,285 | 2,319,971 | 852,115 | 968,388 |
| Largest contig [bp] | 8,233,435 | 6,580,969 | 3,281,684 | 5,352,942 |
| Repeat content | 63.98% | 64.23% | 29.4% | 29.67% |

**Table 2.** BUSCO results of *Prymnesium parvum* ODER1 genome annotation for separated haplotypes and merged haplotypes. For comparison, the BUSCO analysis was run also on the *P. parvum* 12B1 annotation. Note that complete BUSCO values of greater than 90% are hard to obtain for haptophyta, even the currently best haptophyta NCBI reference annotation of *Emiliania huxleyi* reaches only 90.4%.

|  | *P. parvum* ODER1<br>(this study) |  |  | *P. parvum* 12B1<br>(Wisecaver et al.)<sup>16</sup> | *E. huxleyi*<br>(NCBI) |
| --- | --- | --- | --- | --- | --- |
| Haplotype | hap1 | hap2 | hap1 + hap2 | collapsed | collapsed |
| Complete BUSCOs | 88.12% | 89.77% | 90.10% | 84.2% | 90.4% |
| Complete single copy | 83.17% | 86.14% | 3.30% | 77.6% | 89.4% |
| Complete duplicated | 4.95% | 3.63% | 86.80% | 6.6% | 1.0% |
| Fragmented | 3.96% | 3.30% | 3.63% | 3.6% | 4.0% |
| Missing | 7.92% | 6.93% | 6.27% | 12.2% | 5.6% |

Dot plots to compare assemblies of strain 12B1 (type A) and ODER1 (type B) show a conserved collinearity/synteny between the 1n = 34 Hi-C supported chromosomes in both types A and B. This excludes that size differences are due to large-scale duplications or polyploidization (Figure 3A). Regarding structural variation, only a few inter-chromosomal rearrangements become visible, while intra-chromosomal rearrangements occur more often. This is especially true for chromosome1 of ODER1, which matches Scaf2 of strain 12B1 (Figure 3A).

**Figure 3.**
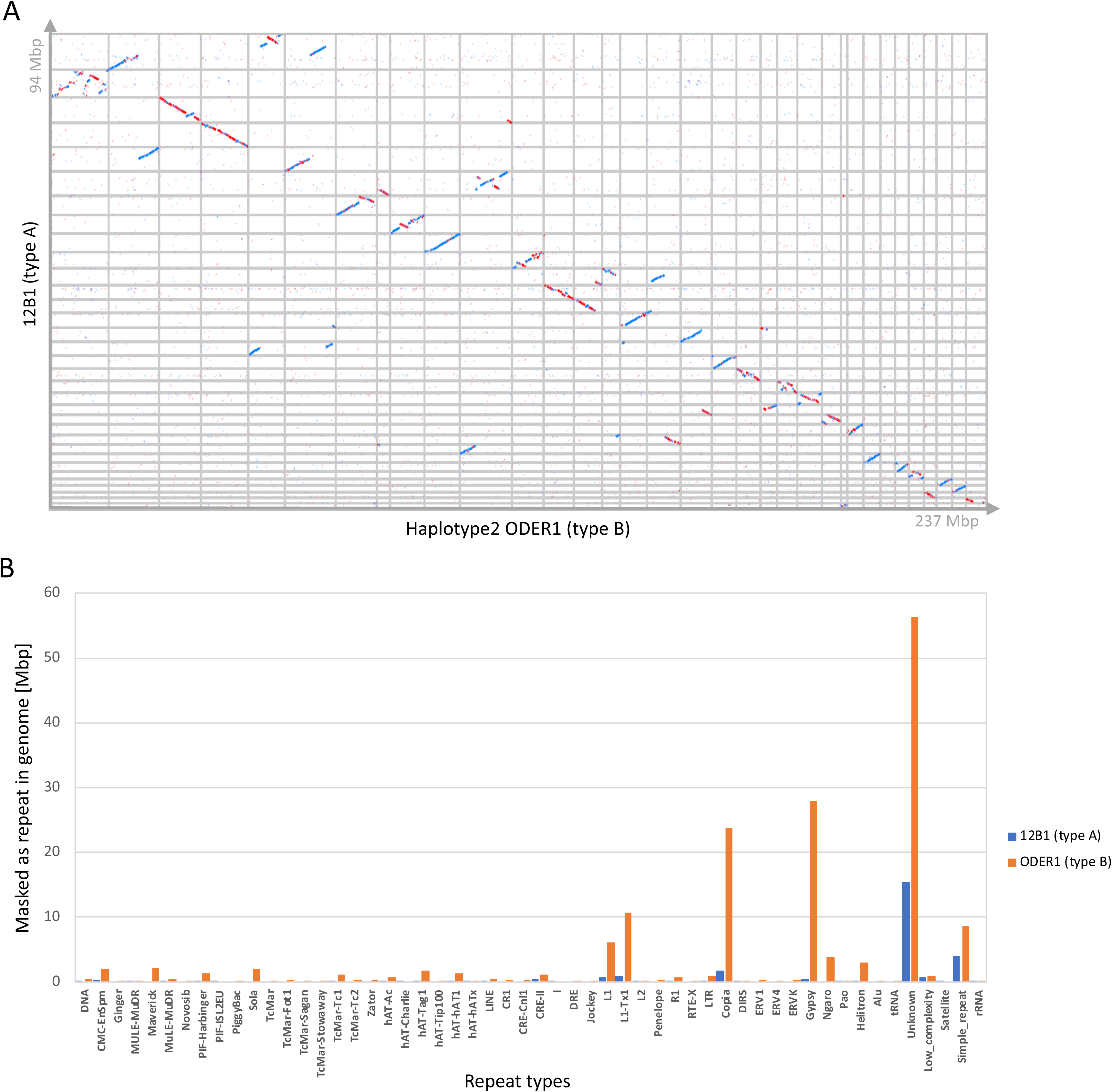
Haploid genome size expansion between *Prymnesium parvum* types A and B. (A) Dotplot comparison between assembled chromosomes of diploid *P. parvum* ODER1 (type B, X-axis) and diploid *P. parvum* 12B1 (type A; Y axis) shows enormous though relatively evenly-distributed size expansion of all chromosomes in the ODER1 strain and some structural rearrangements (blue = forward strand; red = reverse complement strand). (B) Expansion of several repeat classes (L1, Copia, Gypsy, Unknown) explains major genome size differences between the two type A and B strains.

*De novo* repeat annotation showed that the genome size augmentation of ODER1 is due to expansion of repetitive elements and not due to differences in gene content. Repeat content in *P. parvum* ODER1 is 64% compared to 29–30% in both type A reference genomes. Especially, Gypsy- and Copia-like retrotransposons have expanded in *P. parvum* ODER1 compared to *P. parvum* 12B1 (Figure 3B). Complete open reading frames (ORFs) of retroviral genes can be predicted on many annotated repeat elements, suggesting their recent activity. Recent evolutionary activity is also supported by the detection of haplotype-specific retroelement integrations.

For comparisons between type B strains, only short reads and assemblies thereof were available^16^ and due to the high repeat content we found these assemblies highly fragmented, which hindered comparison on the chromosomal-level. Interestingly, K-0081 that has been described as a tetraploid^16^ is the closest relative of ODER1 in the chloroplast-genome based phylogeny. We mapped the available short reads of K-0081 to the ODER1 assembly and found low divergence between the genomes and an allele frequency spectrum (Figure 4A) with a peak at around 33% of heterozygous variant read coverage in K-0081, which hints more at triploidy than at tetraploidy, while the allele frequency plot for ODER1 clearly supported a 50% peak of heterozygous variant reads as expected for a diploid organism. To further investigate the DNA content difference and to assess ploidy level, we performed Propidium Iodide (PI) flow cytometry measurements of strains ODER1 and K-0081. The DNA content of strain ODER1 is 0.55 ± 0.01 pg (534 Mbp), supporting diploidy, while strain K-0081 has a DNA content of 0.77 ± 0.02 pg (756 Mbp), supporting triploidy (Table 3). The DNA content difference between strains was further verified by simultaneous analysis (Figure 4B).

**Figure 4.**
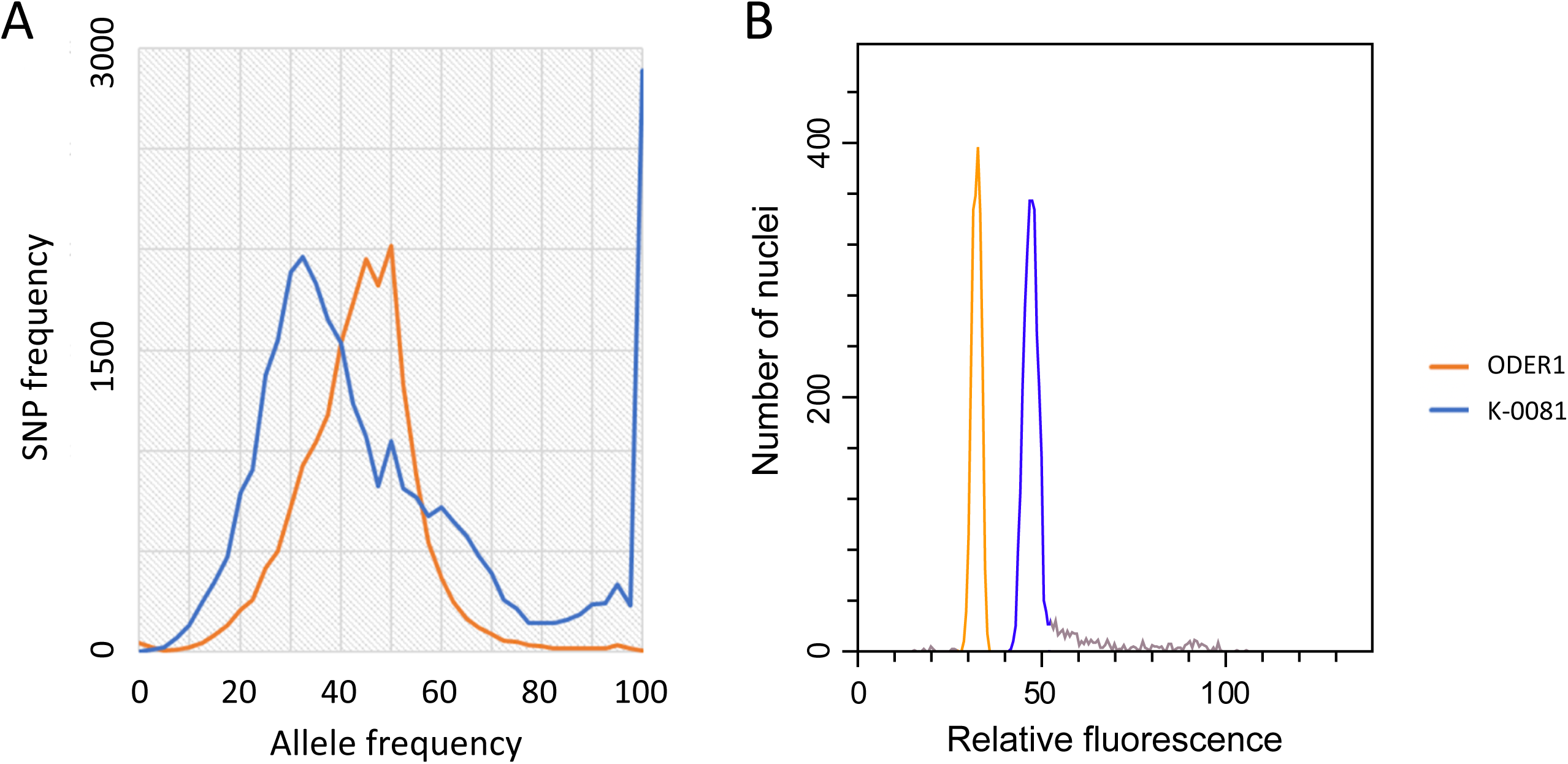
Ploidy analysis of closely related strains ODER1 and K-0081. (A) As expected for a diploid organism, the ODER1 strain showed a peak at 50% allele frequency, while the K-0081 strain had a peak at 33% and a shoulder around 60%, hinting more at triploidy than at tetraploidy. Overall, the number of homozygous variants in K-0081 was low (28,884 variants with 100% allele frequency and coverage >=20x) underlining the very close relationship of both strains. (B) Simultaneous flow cytometric analysis of the type B strains ODER1 and K- 0081. The relative fluorescence of propidium iodide-stained nuclei shows ploidy-level difference between the strains. The resulting DNA content of strain ODER1 (0.55 pg) corresponds to diploidy, while DNA content of strain K-0081 (0.77 pg) corresponds to triploidy.

**Table 3.** Summary statistics of the flow cytometry data of *Prymnesium parvum* strains ODER1 and K-0081.

|  | Replicate | Sample/standard<br>ratio of relative<br>fluorescence | Absolute<br>DNA content<br>(pg) | Absolute DNA<br>content (Mbp) | Sample<br>CV* (%) | Standard CV*<br>(%) |
| --- | --- | --- | --- | --- | --- | --- |
| ODER1 | 1 | 0.668 | 0.548 | 536 | 4.62 | 4.85 |
|  | 2 | 0.648 | 0.532 | 520 | 4.33 | 4.75 |
|  | 3 | 0.682 | 0.559 | 547 | 4.40 | 4.43 |
| K-0081 | 1 | 0.309 | 0.800 | 783 | 4.16 | 3.7 |
|  | 2 | 0.293 | 0.759 | 742 | 4.76 | 3.37 |
|  | 3 | 0.294 | 0.760 | 743 | 4.58 | 3.50 |

\*CV = coefficient of variation of nuclear fluorescence intensity.

The microfibrils of the cell-covering scales, which consist of proteins and carbohydrates, showed a radial arrangement on both faces (proximal and distal) in the ODER1 strain (Figure 5A). Only very rarely, a single scale for which a radial pattern could not be unambiguously identified was found. This is in line with the previously postulated hypothesis that *Prymnesium* cells in the diploid stage lack scales with spirally wound microfibrils on the distal face, which frequently occur in haploid stages in addition to the scales with radially arranged microfibrils.^6,8,49,51^ The scales of the triploid K-0081 strain, in contrast, showed both types of microfibril arrangement (Figure 5B), allowing to distinguish ODER1 and K-0081 based on morphological features. The thickness of the inflexed rim of the scales, which cover the cell in two layers, varied in both strains from narrow (outer layer) to wide (inner layer).

**Figure 5.**
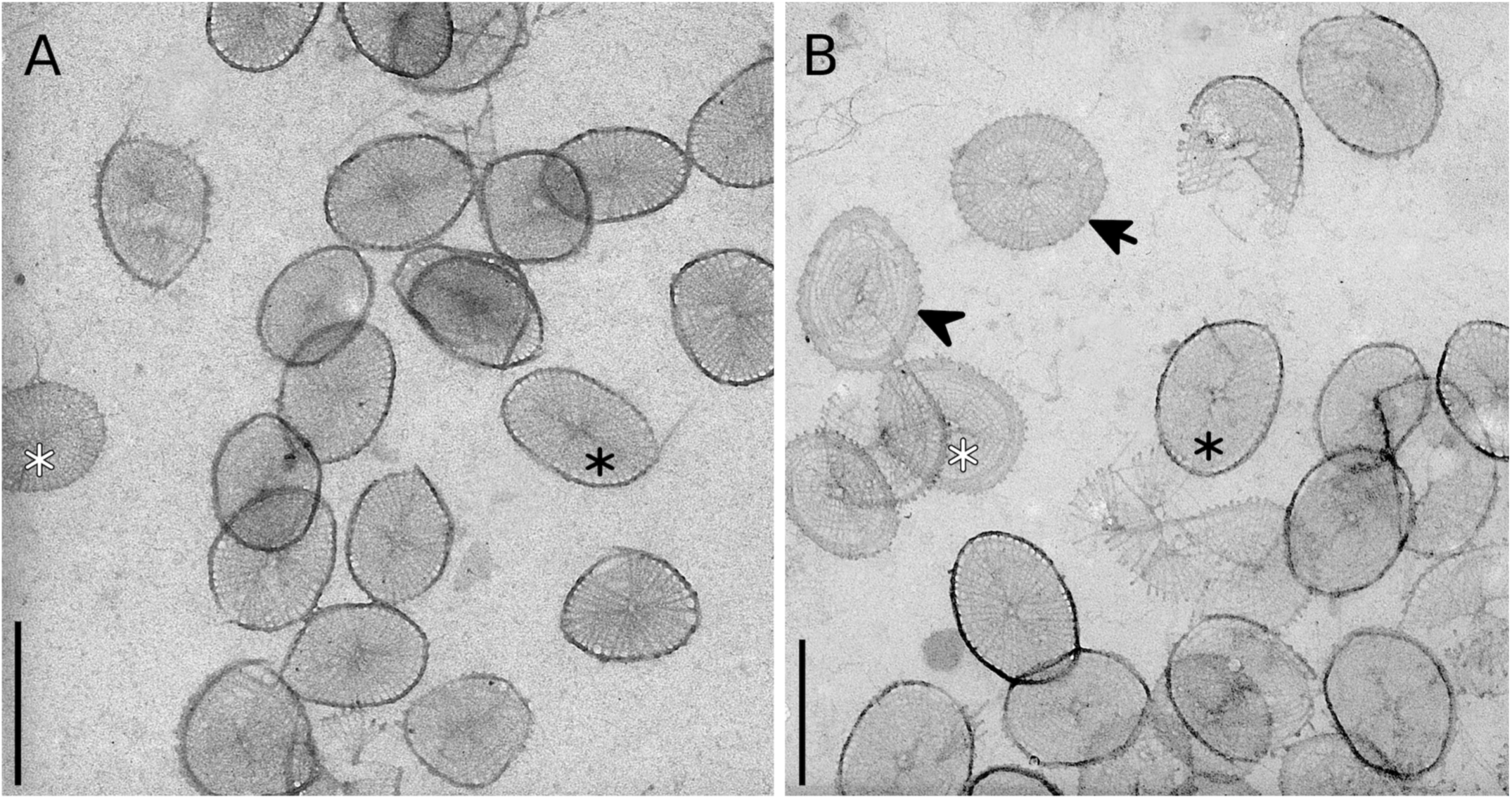
Morphology of *Prymnesium parvum* scales. Transmission electron microscopy (TEM) showing the scales of (A) *P. parvum* strain ODER1 and (B) strain K-0081. Body scales of both layers are shown. Scales of the outer and inner layer are characterized by a narrow (black asterisk) and wide (white asterisk) rim, respectively. Scales of ODER1 show a radial arrangement of microfibrils on both faces whereas those of K-0081 show a radial pattern on the proximal face (arrow) and a spirally wound pattern on the distal face (arrowhead). Scale bars: 400 nm.

### Analyses of polyketide synthases in P. parvum type A and B long-read reference genomes

Polyketide synthases (PKS) are involved in the synthesis of the fish killing toxins, prymnesins, and can be extremely large proteins due to their repeated domain structure. Some PKS identified in *P. parvum* type A genomes hold the ‘world record in protein size’ (>40,000 amino acid residues) and have been named ‘PKZILLA-1’ and ‘PKZILLA-2’.^17^ The domain structure of both enzymes has been related to the prymnesin structure. Due to the presence of large exons, small introns, and repetitiveness, the corresponding genes pose a challenge to most annotation tools and short read transcriptomics. Interestingly, we have found that an *ab initio* annotation tool, namely Genscan,^43^ performed best on the prediction of PKS. This is probably due to the algorithm’s property to produce maximized ORF lengths and caring less for typical exon/intron sizes. Using the two haploid ODER1 assemblies and the other available *P. parvum* type A long-read assemblies (12B1, CCMP3037 and UTEX2797),^15,16^ we were able to predict 26 phylogenetic clades of PKS gene sequences (Figure 6A). While most sequence clades (n = 18) had the corresponding PKS gene present in all genomes examined, we found three clades, where the PKS gene was deleted and one clade, where the PKS gene was tandemly repeated in ODER1 (Figures 6A, B). This underlines divergent evolution of the PKS family among type A and B strains, and is likely to contribute to the genomic basis for the production of structurally different toxic prymnesins A and B. Interestingly, we also found differences in PKS tandem duplications between the haplotype assemblies of ODER1, hinting at very recent evolutionary changes (Figure 6C).

**Figure 6.**
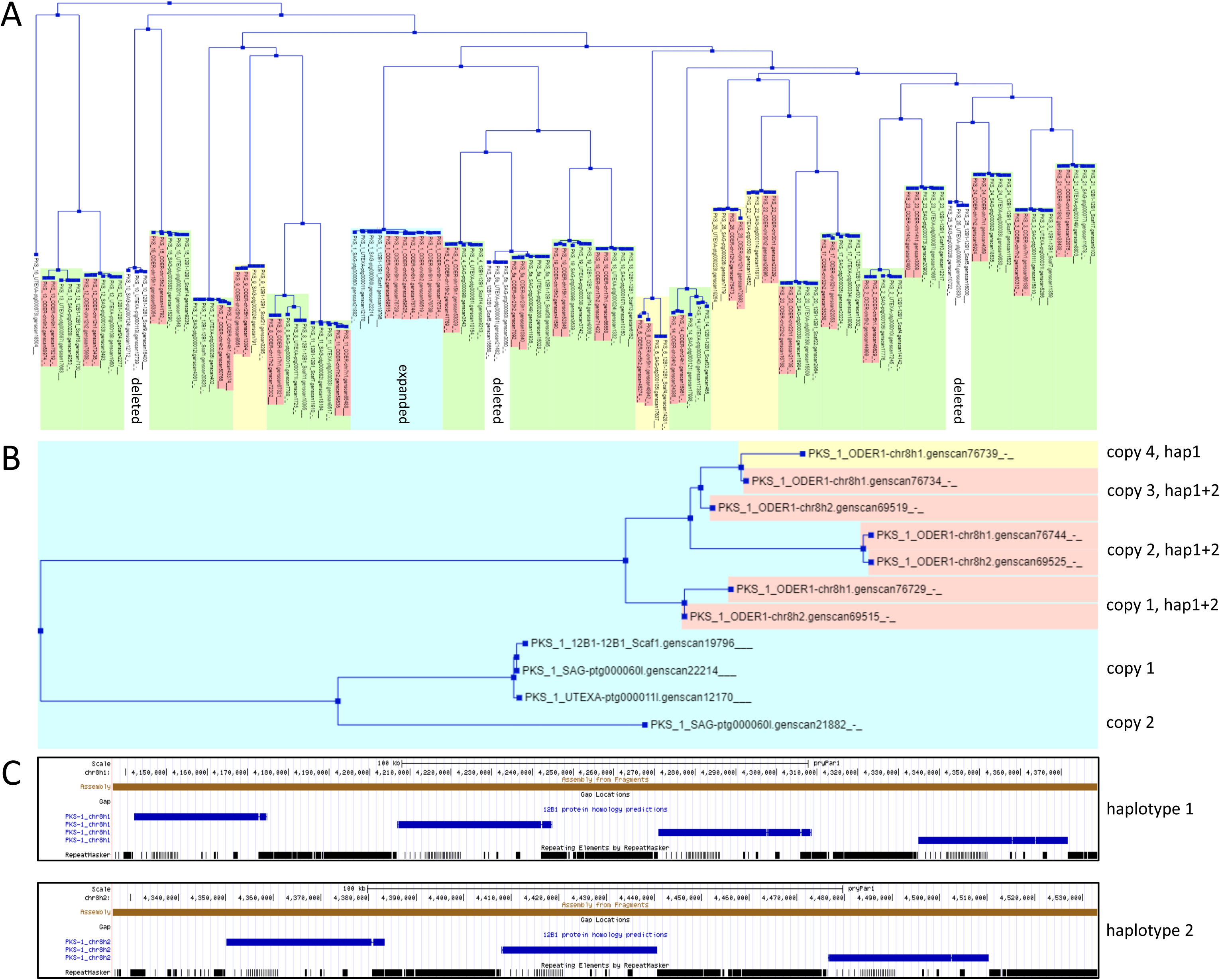
Copy number variation among polyketide synthases in *Prymnesium parvum* types A and B. (A) We identified 26 clades of polyketide synthase (PKS) genes in four *P. parvum* reference genomes (type B: ODER1 (hap1 and hap2, marked in red); type A: 12B1, UTEX2797, CCMP3037). Genes corresponding to 18 clades (green) were found in all four genomes. Four clades (yellow) had missing genes in some of the A-type genomes, which may be due to deletions or assembly issues. Three clades (white) contain genes only in the A-type genomes and none in the *P. parvum* ODER1 genome (neither hap1 nor hap2), suggesting deletion of the genes in the type B ODER1 genome. (B) A single PKS clade (light blue) shows copy number expansion in the ODER1 strain, while type A strains have at most two copies of this PKS gene, the ODER1 strain showed three copies in both haplotypes and even a fourth copy in haplotype 1, which suggests a very recent gene amplification. (C) Plotting these genes along the genomic sequence revealed that they arose by tandem triplication in haplotype 2 and tandem quadruplication in haplotype 1 of *P. parvum* ODER1.

### A deletion in the PKS gene of ODER1 explains structural differences of type A and B prymnesins

During our PKS family analysis a size difference of the largest predicted PKS protein (also referred to as ‘PKZILLA-1’ according to Fallon et al.^17^ between ODER1 and the type A strains became evident, while another large PKS (‘PKZILLA-2’) was conserved (Figure 7). Comparing the corresponding genomic regions between ODER1 and the CCMP3037 assemblies (both from HIFI long-read data assembled without gaps in the regions), revealed a large ODER1-specific deletion (∼20 kbp) and a smaller ODER1 specific duplication (∼4 kbp). Interproscan of the corresponding sequences showed that the deletion removes six KS3_2 domains from ODER1 PKZILLA-1, while the duplication adds one KS3_2 domain (Figures 8A, B). The larger deletion was also found in other type B strains by mapping published short reads against the CCMP3037 assembly and inspecting read coverage (Figure 9). In sum, the five missing KS3_2 domains in type B strain PKZILLA-1 could explain the missing 1,6-dioxadecalin core unit (Figure 7, highlighted in red)^10^ in the structure of the prymnesin B toxins.

**Figure 7.**
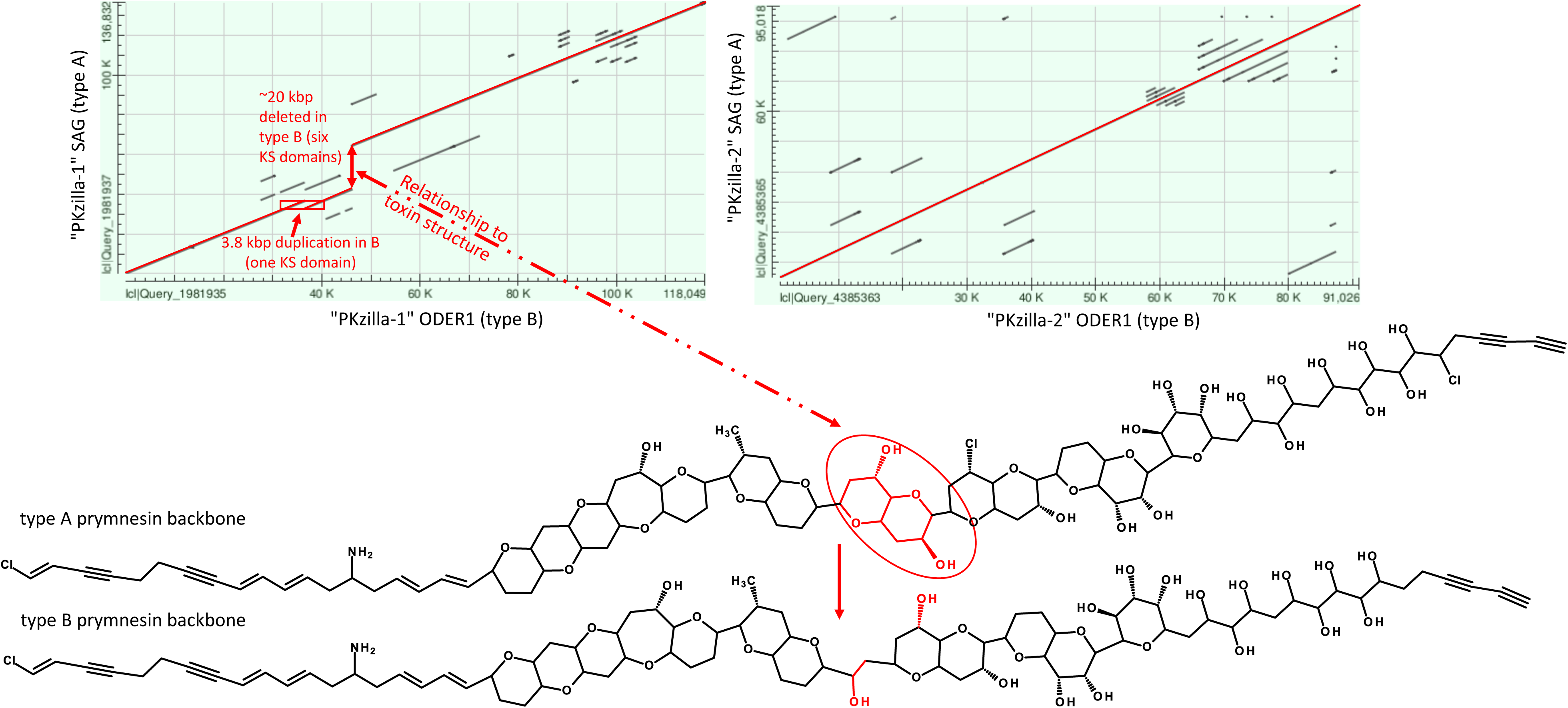
Identification of structural changes in the largest PKS in *Prymnesium parvum* type B versus type A. Structural differences between PKzilla-1 genes in type A and type B strains may correspond to structural changes of the prymnesin toxins produced by these genes. The backbone structure of type A prymnesins according to Igarashi et al.^53^ with three incorporated chlorine atoms (C-1, C- 56, and C85) and those of type B prymesins according to Rasmussen et al.^12^ with one chlorine at position C-1 are displayed. While the type A prymnesin backbone consists of 91 carbons, the type B one has only 85 carbon atoms due to the replacement of one 1,6-dioxadecalin core unit (marked in red) with a short acyclic C2-linkage.

**Figure 8.**
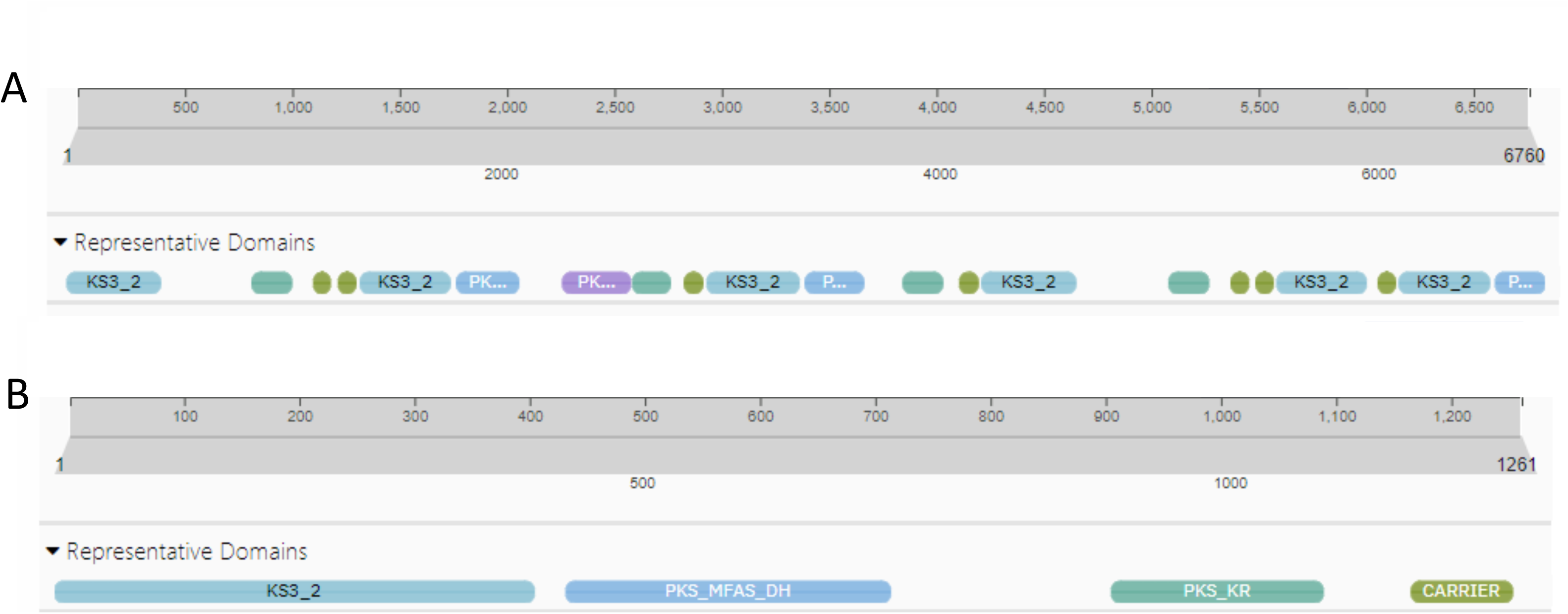
Identification of structural changes in the largest PKS in *Prymnesium parvum* type B versus type A. (A) Six KS3_2 protein domains are deleted in type B (∼20 kbp deletion in A). (B) A single KS3_2 domain is duplicated in type B (3.8 kbp duplication in A).

**Figure 9.**
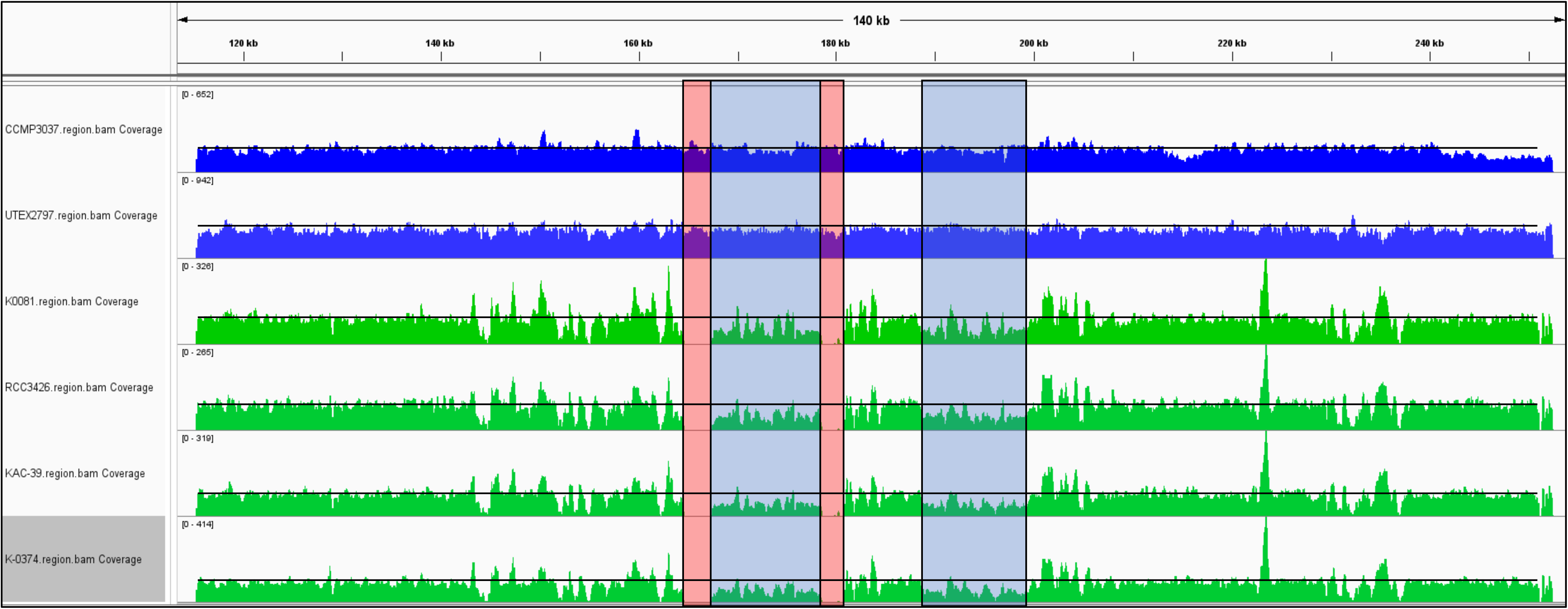
Structural differences of the largest PKS between several type A and B *Prymnesium parvum*. PKZILLA-1 deletion in four additional type B strains (green: K-0081, K-0374, KAC-39, and RCC3426) compared to two type A strains (blue: CMP3037 and UTEX2797) revealed by short read mapping to genomic region of type A PKZILLA-1 (from genome assembly of CCMP3037). Due to the repetitive nature of the PKZILLA-1 gene, large parts of the gene cannot be covered by uniquely mapped reads. Non-uniquely mapped reads are randomly distributed between two larger regions (light blue) that show reduced (∼50%) read-coverage in type B, indicating deletion in type B and duplication in type A. Additionally, borders of the left region do not have any read coverage (red), further supporting the deletion of this part of the gene in type B strains.

## Discussion

Our work exposes in an unprecedented detail the biological agent that by its toxin production mechanistically caused the primarily anthropogenic Oder River disaster in 2022. To our knowledge, this study presents the first reference quality assembly of a type B *P. parvum* and likewise the first haplotype-resolved genome of a haptophyte microalga. Our ODER1 reference genome provides insights into the genetic basis and variability of the toxin production. The chemical structure of the type B toxin backbone differs from type A prymnesins due to the lack of a 1,6-dioxadecalin-core unit^1^ and also the underlying PKS genes show type-specific differences on the level of gene families and gene structures (Figures 6–9) . The structural differences of the largest PKS gene between type A and B result in the gain or loss of several ketide -synthase domains (n = 5) in the respective proteins. This A *vs*. B change in domain number is close to the theoretical counts (n = 3 to 4) that have been predicted based on the number of C-atoms in the prymnesin backbone.^10^ We thus hypothesize that an evolutionary change in the PKS genes, responsible for toxin production, could be the evolutionary basis for a typical B-type toxin. Oder- River-specific prymnesin modifications are characterized by different expressions of certain prymnesins, which might be connected to differences of K-0081 and ODER1 regarding a glycosyltransferase gene (Figure 10).^1^ Whether or not this is relevant for a specific toxicity^18^ is currently studied in the ODER-SO project (https://www.igb-berlin.de/en/oder-so).

**Figure 10.**
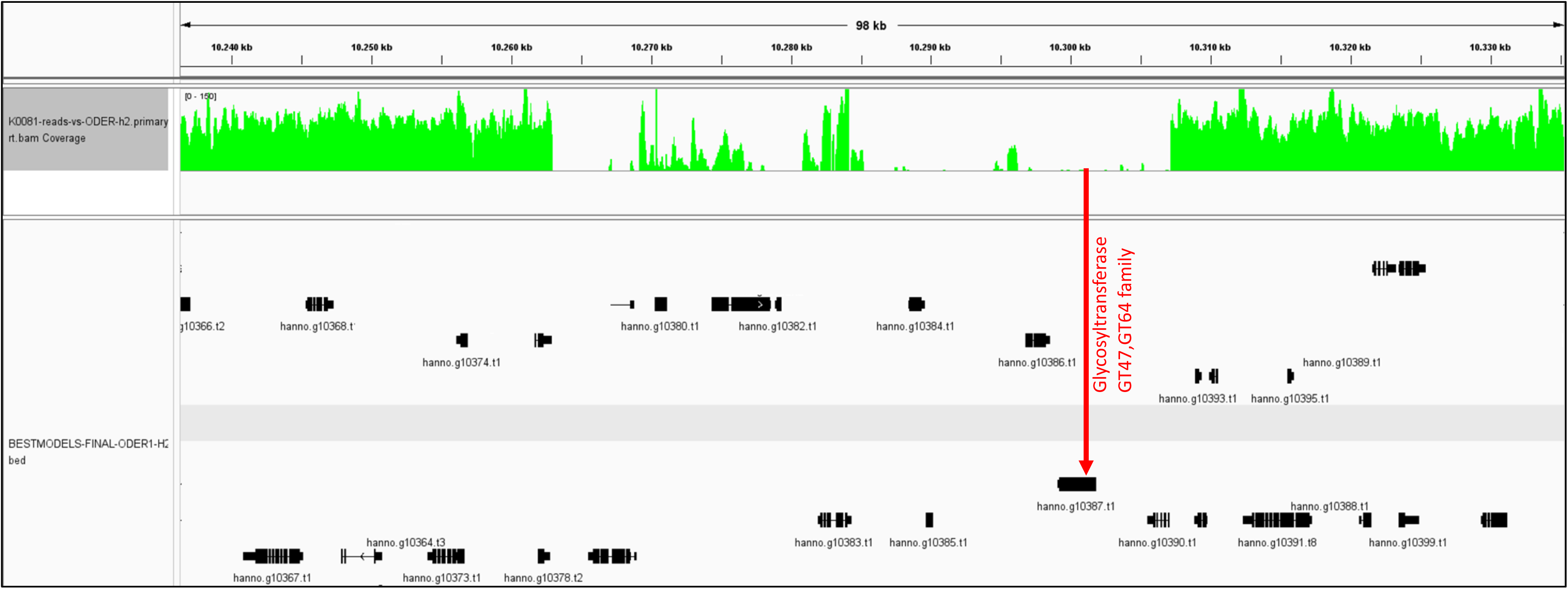
Glycosyltransferase gene present in *Prymnesium parvum* ODER1 but missing in strain K-0081. A glycosyltransferase (CAZy-ID GT47/GT64) in ODER1 is missing in K-0081 as shown by zero short-read coverage obtained by whole genome sequencing (green) of K-0081 when mapped to the ODER1 haplotype 2.

This reference genome further adds to the evidence by Wisecaver et al.^16^ that the hidden, probably ‘species-level’ diversity within *P. parvum* s.l. can only be comprehensively understood using state-of-the-art genomics. In this regard, the ODER1 assembly strongly suggests the expansion of retroelements as a driver of haploid genome size evolution, as earlier concluded from indirect methods (estimated from highly fragmented short-read assemblies).^16^ Their high repeat content renders genome assembly of type B more complex than type A strains and underlines the need for high-accuracy long-read approaches. In the future, repetitive (retro)viral sequences in the genome may help to discover types of potentially infectious viruses in environmental samples that might serve as agents of biological pest control to fight *P. parvum* blooms.^50^

Besides haploid genome size evolution, differences or shifts in ploidy contribute to diversity among *P. parvum* s.l. and complicate the ‘big picture’ to systematize these microalgae. Comparative flow-cytometric analyses as well as our genomic and morphological data clearly show that the ODER1 strain is a diploid form. During their life cycle, *P. parvum* s.l. microalgae alternate between haploid and diploid life stages, each of which can reproduce by asexual mitotic divisions^8^ and may be able to generate blooms. Thus, genotypically highly similar if not identical life forms occur. According to our analyses, the closest known relative of the diploid ODER1 strain is K-0081 from Denmark. However, our flow-cytometric measurements and analysis of SNP allele frequency distributions (Figure 4) revealed K-0081 to be triploid. It remains unclear whether this indicates previously undescribed genetic/genomic plasticity in these algae, or more likely an artifact resulting from nearly four decades of K-0081 cultivation. Of note, K-0081 was previously described as tetraploid, possibly due to an underestimation of its haploid genome size based on short-read sequencing in combination with flow cytometric data that often result in 10–20% larger genome size estimates than sequencing data.^52^

While scale morphology supports diploidy of ODER1, it has not been examined in triploid *P. parvum* s.l. before. Thus, our morphological data neither corroborates nor contradicts the K- 0081 triploidy inferred from flow-cytometric and SNP allele frequency analyses. The triploid K- 0081 exhibits radial and spirally wound microfibril arrangement, so far known only in haploid cells.^6,8,49,51^

Together with the so far existing type A reference genomes, our high-quality genome of type B presents an important basic research contribution for comparative genomic analyses of this globally relevant group of microalgae. Using short-read sequencing, the reference genomes (type C is pending) now allow, when detected in a specific region, taxonomically determining *P. parvum* s.l. with little effort in whole genome detail. This will form the basis for the development of taxon-specific control methods and surveillance of its potential evolutionary adaptation and change. The link between PKS and toxins enables a better understanding of the mechanistic relationships between gene expression, toxin production and ecological/environmental conditions. To understand the interaction of these factors in natural water bodies, growth and toxicity experiments are required to predict *P. parvum* strain-specific blooms and the causal ecological conditions. *Prymnesium parvum* remains present in the entire Oder River after the summer 2022 bloom as documented by us using a molecular-quantification (qPCR) assay. It shows the presence of this microalga (03/2023–02/2024), including major shifts in quantities (Figure 11), suggesting a potential for future bloom threats, when the compound environmental conditions, triggering such a disaster, may be fulfilled.^1^

**Figure 11.**
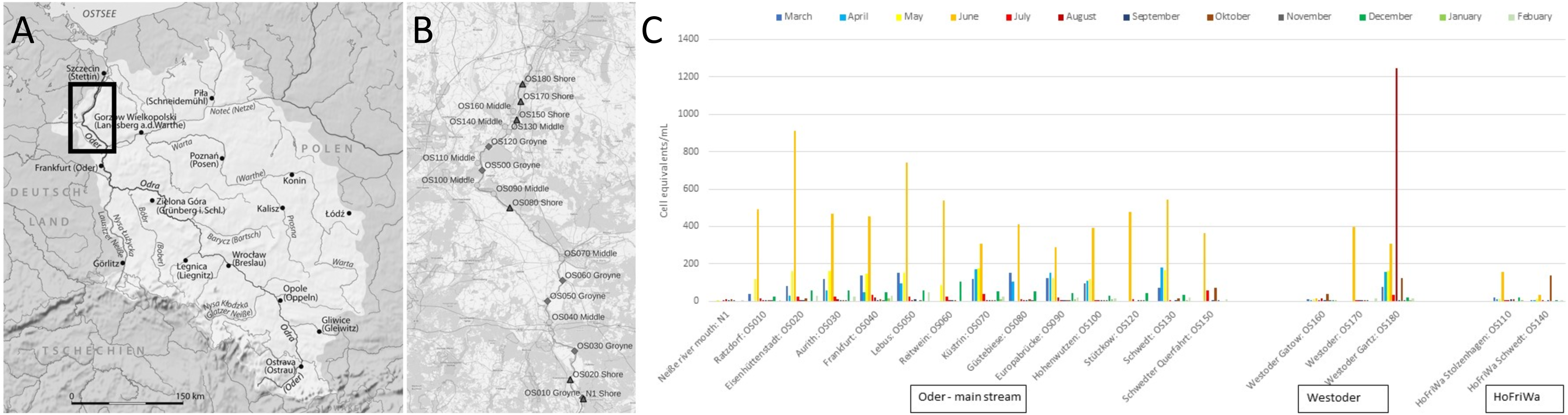
Monitoring with abundances of *Prymnesium parvum* in the Oder River catchment using qPCR quantification. (A) Overview map and (B) sampling sites along the German part of the Oder River. (C) Cell equivalents determined as ITS1 copies/mL shown from March 2023 until February 2024 in the Oder mainstream, the West-Oder River branch, and channel near Hohensaatener- Friedrichsthaler-Wasserstraße (HoFriWa).

## Acknowledgements

We thank colleagues at the Leibniz Institute of Freshwater Ecology and Inland Fisheries (IGB) for help with field and laboratory work, especially Katrin Preuss, Miriam Diemer, Shambhavi Parmar, and Karla Münzer, who raised the *Prymnesium* cultures and prepared samples, and Martin Pusch for supporting our research proposal. We thank Susan Mbedi (Berlin Center for Genomics in Biodiversity Research) for help with sequencing and Renate Radek (Freie Universität Berlin) for kind assistance with electron microscopy. *Prymnesium parvum* strain K-0081 was obtained from the Norwegian Culture Collection of Algae (NORCCA, Oslo); for strains UTEX2797 and RCC1436, we thank Per Juel Hansen (University of Copenhagen). DČ is grateful for the financial support of the Martina Roeselová Memorial Fellowship granted by the IOCB Tech Foundation.

Our study was conducted as part of the project ODER∼SO, financed by the German Federal Agency for Nature Conservation (BfN) with funds from the German Federal Ministry for the Environment, Nature Conservation, Nuclear Safety and Consumer Protection (BMUV).

## Author contributions

Conceptualization, MS, MTM, DKL

Resources, JK, JFHS, MS

Data collection and curation, HK, JS, DČ, EK Software, HK

Formal analysis, HK, JFHS, DČ, SW

Funding acquisition, MS, MTM

Writing – original draft, HK, MS, JFHS, EV, MTM

Writing – review and editing, all authors

## References

1. Köhler, J., Varga, E., Spahr, S., Gessner, J., Stelzer, K., Brandt, G., Mahecha, M.D., Kraemer, G., Pusch, M., Wolter, C., et al. (2023). Unpredicted ecosystem response to compound human impacts in a European river. Res Sq.

2. Szlauer-Łukaszewska, A., Ławicki, Ł., Engel, J., Drewniak, E., Ciężak, K., and Marchowski, D. (2024). Quantifying a mass mortality event in freshwater wildlife within the Lower Odra River: Insights from a large European river. Sci Total Environ 907, 167898.

3. Sobieraj, J., and Metelski, D. (2023). Insights into toxic Prymnesium parvum blooms as a cause of the ecological disaster on the Odra River. Toxins (Basel) 15.

4. Sproston, N.G. (1946). Fish mortality due to a brown flagellate. Nature 158, 70–71.

5. 5. Our Molecular Biology Correspondent (1967). Polymerase activity and dissociation. Nature *216*, 852–853.

6. Green, J.C., Hibberd, D.J., and Pienaar, R.N. (1982). The taxonomy of Prymnesium (Prymnesiophyceae) including a description of a new cosmopolitan species, P. patellifera sp. nov., and further observations on P. parvum N. carter. Br Phycol J 17, 363–382.

7. Larsen, A., and Medlin, L.K. (1997). Inter- and intraspecific genetic variation in twelve Prymaesium (haptophyceae) clones. J Phycol 33, 1007–1015.

8. Gastineau, R., Davidovich, N.A., Hallegraeff, G., Probert, I., and Mouget, J.-L. (2014). Reproduction in microalgae. In Reproductive Biology of Plants, K. G. Ramawa, J.-M. Mérillon, and K. R. Shivanna, eds. (CRC Press), pp. 1–28.

10. Lutz-Carrillo, D.J., Southard, G.M., and Fries, L.T. (2010). Global genetic relationships among isolates of golden alga (Prymnesium parvum). JAWRA J Am Water Resour Assoc 46, 24–32.

11. Binzer, S.B., Svenssen, D.K., Daugbjerg, N., Alves-de-Souza, C., Pinto, E., Hansen, P.J., Larsen, T.O., and Varga, E. (2019). A-, B- and C-type prymnesins are clade specific compounds and chemotaxonomic markers in Prymnesium parvum. Harmful Algae 81, 10–17.

12. Driscoll, W.W., Wisecaver, J.H., Hackett, J.D., Espinosa, N.J., Padway, J., Engers, J.E., and Bower, J.A. (2023). Behavioural differences underlie toxicity and predation variation in blooms of Prymnesium parvum. Ecol Lett 26, 677–691.

13. Rasmussen, S.A., Meier, S., Andersen, N.G., Blossom, H.E., Duus, J.Ø., Nielsen, K.F., Hansen, P.J., and Larsen, T.O. (2016). Chemodiversity of ladder-frame prymnesin polyethers in Prymnesium parvum. J Nat Prod 79, 2250–2256.

14. Koid, A.E., Liu, Z., Terrado, R., Jones, A.C., Caron, D.A., and Heidelberg, K.B. (2014). Comparative transcriptome analysis of four prymnesiophyte algae. PLoS One 9, e97801.

15. Anestis, K., Kohli, G.S., Wohlrab, S., Varga, E., Larsen, T.O., Hansen, P.J., and John, U. (2021). Polyketide synthase genes and molecular trade-offs in the ichthyotoxic species Prymnesium parvum. Sci Total Environ 795, 148878.

16. Jian, J., Wu, Z., Silva-Núñez, A., Li, X., Zheng, X., Luo, B., Liu, Y., Fang, X., Workman, C.T., Larsen, T.O., et al. (2024). Long-read genome sequencing provides novel insights into the harmful algal bloom species Prymnesium parvum. Sci Total Environ 908, 168042.

17. Wisecaver, J.H., Auber, R.P., Pendleton, A.L., Watervoort, N.F., Fallon, T.R., Riedling, O.L., Manning, S.R., Moore, B.S., and Driscoll, W.W. (2023). Extreme genome diversity and cryptic speciation in a harmful algal-bloom-forming eukaryote. Curr Biol 33, 2246–2259.e8.

18. Fallon, T.R., Shende, V. V, Wierzbicki, I.H., Auber, R.P., Gonzalez, D.J., Wisecaver, J.H., and Moore, B.S. (2024). Giant polyketide synthase enzymes biosynthesize a giant marine polyether biotoxin. bioRxiv, 2024.01.29.577497.

18. Varga, E., Prause, H.-C., Riepl, M., Hochmayr, N., Berk, D., Attakpah, E., Kiss, E., Medić, N., Del Favero, G., Larsen, T.O., et al. (2024). Cytotoxicity of Prymnesium parvum extracts and prymnesin analogs on epithelial fish gill cells RTgill-W1 and the human colon cell line HCEC-1CT. Arch Toxicol 98, 999–1014.

19. Auber, R., and Wisecaver, J.H. (2019). Algal nuclei isolation for nanopore sequencing of HMW DNA V.3.

20. Ruan, J., and Li, H. (2020). Fast and accurate long-read assembly with wtdbg2. Nat Methods 17, 155–158.

21. Kolmogorov, M., Yuan, J., Lin, Y., and Pevzner, P.A. (2019). Assembly of long, error-prone reads using repeat graphs. Nat Biotechnol 37, 540–546.

22. Li, H. (2018). Minimap2: pairwise alignment for nucleotide sequences. Bioinformatics 34, 3094–3100.

23. Frith, M.C., and Kawaguchi, R. (2015). Split-alignment of genomes finds orthologies more accurately. Genome Biol 16, 106.

24. Blanchette, M., Kent, W.J., Riemer, C., Elnitski, L., Smit, A.F.A., Roskin, K.M., Baertsch, R., Rosenbloom, K., Clawson, H., Green, E.D., et al. (2004). Aligning multiple genomic sequences with the threaded blockset aligner. Genome Res 14, 708–715.

25. Minh, B.Q., Schmidt, H.A., Chernomor, O., Schrempf, D., Woodhams, M.D., von Haeseler, A., and Lanfear, R. (2020). IQ-TREE 2: new models and efficient methods for phylogenetic inference in the genomic era. Mol Biol Evol 37, 1530–1534.

26. Hoang, D.T., Chernomor, O., Von Haeseler, A., Minh, B.Q., and Vinh, L.S. (2018). UFBoot2: improving the ultrafast bootstrap approximation. Mol Biol Evol 35, 518–522.

27. Guindon, S., Dufayard, J.F., Lefort, V., Anisimova, M., Hordijk, W., and Gascuel, O. (2010). New algorithms and methods to estimate maximum-likelihood phylogenies: assessing the performance of PhyML 3.0. Syst Biol 59, 307–321.

28. Cheng, H., Concepcion, G.T., Feng, X., Zhang, H., and Li, H. (2021). Haplotype-resolved de novo assembly using phased assembly graphs with hifiasm. Nat Methods 18, 170–175.

29. Zhang, H., Song, L., Wang, X., Cheng, H., Wang, C., Meyer, C.A., Liu, T., Tang, M., Aluru, S., Yue, F., et al. (2021). Fast alignment and preprocessing of chromatin profiles with Chromap. Nat Commun 12, 6566.

30. Zhou, C., McCarthy, S.A., and Durbin, R. (2023). YaHS: yet another Hi-C scaffolding tool. Bioinformatics 39.

31. Durand, N.C., Robinson, J.T., Shamim, M.S., Machol, I., Mesirov, J.P., Lander, E.S., and Aiden, E.L. (2016). Juicebox provides a visualization system for Hi-C contact maps with unlimited zoom. Cell Syst 3, 99–101.

32. Li, H. (2021). New strategies to improve minimap2 alignment accuracy. Bioinformatics 37, 4572– 4574.

33. Li, H. (2016). Minimap and miniasm: fast mapping and de novo assembly for noisy long sequences. Bioinformatics 32, 2103–2110.

34. Bolger, A.M., Lohse, M., and Usadel, B. (2014). Trimmomatic: a flexible trimmer for Illumina sequence data. Bioinformatics 30, 2114–2120.

35. Haas, B.J., Papanicolaou, A., Yassour, M., Grabherr, M., Blood, P.D., Bowden, J., Couger, M.B., Eccles, D., Li, B., Lieber, M., et al. (2013). De novo transcript sequence reconstruction from RNA- seq using the Trinity platform for reference generation and analysis. Nat Protoc 8, 1494–1512.

36. Flynn, J.M., Hubley, R., Goubert, C., Rosen, J., Clark, A.G., Feschotte, C., and Smit, A.F. (2020). RepeatModeler2 for automated genomic discovery of transposable element families. Proc Natl Acad Sci 117, 9451–9457.

37. Li, H. (2023). Protein-to-genome alignment with miniprot. Bioinformatics 39, btad014.

38. Pertea, M., Pertea, G.M., Antonescu, C.M., Chang, T.-C., Mendell, J.T., and Salzberg, S.L. (2015). StringTie enables improved reconstruction of a transcriptome from RNA-seq reads. Nat Biotechnol 33, 290–295.

39. Niknafs, Y.S., Pandian, B., Iyer, H.K., Chinnaiyan, A.M., and Iyer, M.K. (2017). TACO produces robust multisample transcriptome assemblies from RNA-seq. Nat Methods 14, 68–70.

40. Cantalapiedra, C.P., Hernández-Plaza, A., Letunic, I., Bork, P., and Huerta-Cepas, J. (2021). eggNOG-mapper v2: functional annotation, orthology assignments, and domain prediction at the metagenomic scale. Mol Biol Evol Evol 38, 5825–5829.

41. Kiełbasa, S.M., Wan, R., Sato, K., Horton, P., and Frith, M.C. (2011). Adaptive seeds tame genomic sequence comparison. Genome Res 21, 487–493.

42. Simão, F.A., Waterhouse, R.M., Ioannidis, P., Kriventseva, E. V., and Zdobnov, E.M. (2015). BUSCO: assessing genome assembly and annotation completeness with single-copy orthologs. Bioinformatics 31, 3210–3212.

43. Burge, C., and Karlin, S. (1997). Prediction of complete gene structures in human genomic DNA. J Mol Biol 268, 78–94.

44. Dpooležel, J., Binarová, P., and Lcretti, S. (1989). Analysis of nuclear DNA content in plant cells by flow cytometry. Biol Plant 31, 113–120.

45. Temsch, E.M., Greilhuber, J., and Krisai, R. (2010). Genome size in liverworts. Preslia 82, 63–80.

46. Veselý, P., Bureš, P., Šmarda, P., and Pavlíček, T. (2012). Genome size and DNA base composition of geophytes: the mirror of phenology and ecology? Ann Bot 109, 65–75.

47. Doležel, J., and Bartoš, J. (2005). Plant DNA flow cytometry and estimation of nuclear genome size. Ann Bot 95, 99–110.

48. Zheng, Z., Li, S., Su, J., Leung, A.W.-S., Lam, T.-W., and Luo, R. (2022). Symphonizing pileup and full-alignment for deep learning-based long-read variant calling. Nat Comput Sci 2, 797–803.

49. Larsen, A., and Edvardsen, B. (1998). Relative ploidy levels in Prymnesium parvum and P. patelliferum (Haptophyta) analyzed by flow cytometry. Phycologia 37, 412–424.

50. Wagstaff, B.A., Vladu, I.C., Barclay, J.E., Schroeder, D.C., Malin, G., and Field, R.A. (2017). Isolation and characterization of a double stranded DNA megavirus infecting the toxin-producing haptophyte Prymnesium parvum. Viruses 9.

51. Eikrem, W., Medlin, L.K., Henderiks, J., Rokitta, S., Rost, B., Probert, I., Throndsen, J., and Edvardsen, B. (2017). Haptophyta BT - Handbook of the Protists. In, J. M. Archibald, A. G. B. Simpson, and C. H. Slamovits, eds. (Springer International Publishing), pp. 893–953.

52. Kasai, F., O’Brien, P.C.M., and Ferguson-Smith, M.A. (2013). Afrotheria genome; overestimation of genome size and distinct chromosome GC content revealed by flow karyotyping. Genomics 102, 468–471.

53. Igarashi, T., Satake, M., and Yasumoto, T. (1996). Prymnesin-2: a potent ichthyotoxic and hemolytic glycoside isolated from the red tide alga Prymnesium parvum. J Am Chem Soc 118, 479–480.

